# Nir1-Nir2 Heterodimerization Expands the Sensitivity and Dynamic Range of the Phosphatidylinositol Cycle

**DOI:** 10.64898/2026.06.03.729928

**Authors:** Taylor A. Rahn, Wei-Ting Li, Wan-Ru Lee, Lingshuang Wu, Michael V. Airola, Jen Liou

## Abstract

The phosphatidylinositol (PI) cycle sustains phosphoinositide-calcium signaling by replenishing plasma membrane phosphatidylinositol 4,5-bisphosphate (PIP2) consumed during receptor activation. Recruitment of the lipid transfer protein Nir2 to endoplasmic reticulum (ER)-plasma membrane (PM) contact sites (also known as ER-PM junctions) is central to this process, enabling transfer of PI from the ER to the PM for PIP2 resynthesis. Although the disease-associated Nir2 paralog Nir1 is essential for Nir2 recruitment under physiological conditions, the molecular basis of this regulation has remained unresolved. Here, we identify a conserved Nir Dimerization (NirD) domain in both Nir1 and Nir2 and determine the crystal structures. We show that NirD mediates preferential Nir1-Nir2 heterodimerization, which promotes Nir2 recruitment to ER-PM junctions and enhances PIP2 replenishment in stimulated cells. Furthermore, Nir1-Nir2 heterodimerization confers stimulus-strength-dependent, graded recruitment of Nir2, thereby broadening the sensitivity and dynamic range of PI cycle activity. Together, our findings uncover the structural mechanism underlying Nir1-dependent regulation of Nir2 and reveal paralog heterodimerization as a key strategy for scaling lipid transport at membrane contact sites to signaling demand, ensuring robust phosphoinositide homeostasis.

## INTRODUCTION

Phosphatidylinositol 4,5-bisphosphate (PIP_2_) is an anionic phospholipid enriched in the inner leaflet of the plasma membrane (PM) ^1^. Maintenance of PM PIP_2_ levels is essential for numerous cellular processes, including actin cytoskeleton remodeling, endocytosis and exocytosis, and ion transport ^2^. PIP_2_ also serves as a substrate for phospholipase C (PLC), which is activated downstream of cell surface receptors and hydrolyzes PIP_2_ to generate inositol 1,4,5-trisphosphate (IP_3_) and diacylglycerol (DAG), thereby triggering intracellular calcium signaling and protein kinase C activation. PLC-mediated PIP_2_ consumption necessitates rapid and efficient lipid replenishment through the phosphatidylinositol (PI) cycle to sustain signaling and preserve cellular homeostasis in response to a broad range of extracellular stimuli, including hormones, growth factors, neurotransmitters ^3,4^.

The PI cycle depends on lipid transfer proteins that shuttle phosphatidic acid (PA), generated from DAG at the PM, to the endoplasmic reticulum (ER) for PI synthesis, and deliver PI back to the PM for PIP_2_ resynthesis ^5^. This lipid exchange is mediated by Class IIA PI transfer proteins (PITPs) Nir2 and Nir3 (also known as PITPNM1 and PITPNM2), which operate at ER-PM junctions, specialized membrane contact sites where the ER and the PM are separated by less than 30 nanometers ^6–8^. Loss of Nir2 and Nir3 disrupts PIP_2_ regeneration following receptor stimulation ^6^, and impairs multiple PIP_2_-dependent processes, including phagocytosis ^9^, cytokinesis ^10^, cell motility ^11^, and lymphocyte development ^12^.

Nir1 (also known as PITPNM3) is the third member of this protein family but lacks the N-terminal PITP domain present in Nir2 and Nir3. In contrast to Nir2 and Nir3, Nir1 constitutively localizes to ER-PM junctions and drives the recruitment of Nir2 ^13^. Consistent with its *Drosophila* homolog RdgB, Nir1 also plays essential roles in the retina, and its mutation is linked to autosomal cone dystrophy ^14–17^. Despite lacking PI transfer activity, Nir1 shares a conserved domain architecture with Nir2 and Nir3, including: 1) an FFAT motif that binds ER-resident VAP proteins ^18^; 2) a C-terminal LNS2 domain, a pseudoenzyme that binds PA rather than hydrolyzing it ^19^, and is required for PM recruitment in response to elevated PA levels ^6,13^; and 3) a conserved DDHD region of previously unknown function, homologous to the catalytic domains of the DDHD1 and DDHD2 lipases ^20,21^.

Here, we establish that the DDHD region constitutes part of a larger structural module, which we define as the Nir dimerization (NirD) domain. We determine experimental structures of the NirD domain and show that the DDHD region has been repurposed to drive Nir dimerization while losing the enzymatic activity characteristic of DDHD1 and DDHD2 lipases. Using rationally designed mutations, we disrupt NirD-mediated dimerization both *in vitro* and in cells and demonstrate that this domain is required for Nir1-dependent recruitment of Nir2 to ER-PM junctions. We further show that Nir1-Nir2 heterodimerization through the NirD domain enhances PIP_2_ replenishment following receptor stimulation. Finally, we demonstrate that dimerization with Nir1 enables graded recruitment of Nir2 in proportion to stimulus intensities, thereby enhancing the robustness of the PI cycle. Together, these findings define a molecular mechanism that enables precise and scalable control of PIP_2_ homeostasis across a wide range of external inputs.

## RESULTS

### Structural predictions of Nir2

Examination of an AlphaFold structural prediction of Nir2 suggested the annotated DDHD domain is not an autonomous structural unit but instead forms part of a larger globular domain of previously unrecognized function. This putative domain spans residues 420-887 between the FFAT motif and LNS2 domain and encompasses the entire DDHD region (residues 686- 880) **(Fig. 1A)**. These observations prompted us to determine whether this predicted structure constitutes a bona fide domain and to define its role in Nir2 function.

**Fig. 1.**
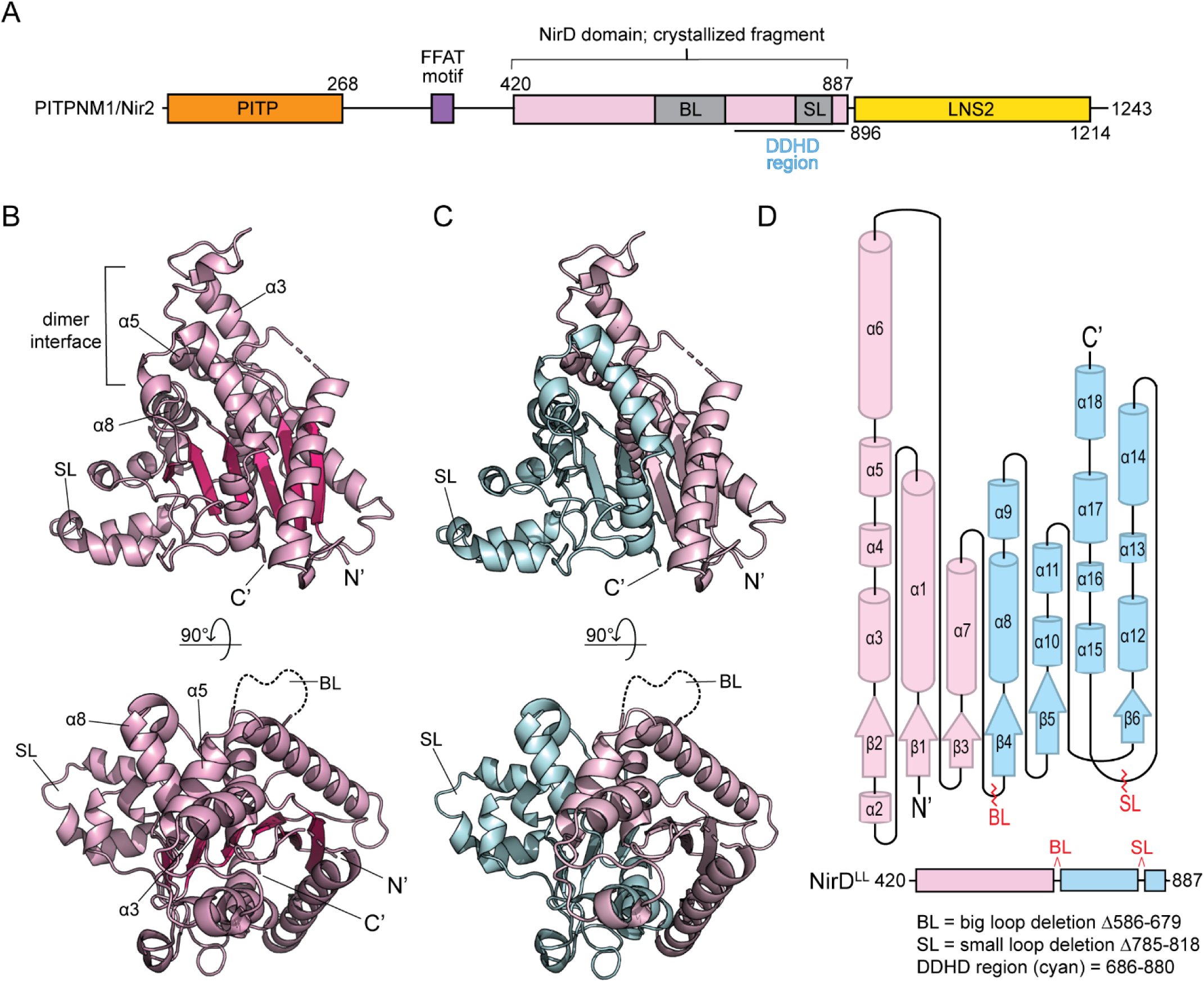
Crystal structure of the Nir2 NirD domain. **(A)** Domain architecture of Nir2 depicting the domain constraints of the NirD domain and the crystallized fragment with the big loop (BL) and small loop (SL) removed. PITP, phosphatidylinositol transfer protein domain; NirD, Nir dimerization domain; LNS2, Lipin/Ned1/Smp2 domain. **(B)** Overall structure of the Nir2-NirD^LL^ domain. Front view (top) shows the dimer interacting regions that involve hydrophobic interactions between the α3, α5, and α8 helices, which contain mainly non-polar residues and exclude water from the interface. The locations of the big loop (BL) and small loop (SL) are highlighted. **(C)** Structure of the Nir2-NirD^LL^ domain shown in identical views as in B, with the DDHD region highlighted. The DDHD region (cyan, residues 686-880) co-folds with the remaining sequence (pink) to form the complete domain. **(D)** Two-dimensional topological map of the secondary structure elements in Nir2-NirD^LL^. The loop deletion regions are labeled in the topological map and the corresponding domain architecture (bottom).

### Structure of the Nir2 NirD domain

We first expressed and purified a recombinant Nir2 fragment corresponding to the predicted domain identified by structural modeling. As discussed below, this region is referred to as the Nir Dimerization (NirD) domain. Despite extensive efforts, we were unable to crystallize the full-length NirD^WT^ domain, likely due to the presence of two predicted intrinsically disordered loops. To facilitate crystallization, we generated a modified construct containing internal deletions of a big loop (BL; residues 586-679) and a small loop (SL; residues 785-818) **(Fig. 1A)**, herein referred to as NirD^LL^ (LL = “loopless”). Nir2-NirD^LL^ successfully crystallized, and the structure was refined to 2.45 Å **(Table 1)**. Of note, while this study was in progress, a similar structure was reported for this region of Nir2 ^22^.

**Table 1.** Data Collection and Refinement Statistics.

|  | **Nir2 NirD<sup>LL</sup>** | **Nir1 NirD<sup>LL</sup>** |
| --- | --- | --- |
| **PDB ID** | 9C37 | 9C38 |
| **Wavelength (Å)** | 0.9793 | 0.9793 |
| **Resolution range (Å)** | 40.92 - 2.45<br>(2.54 - 2.45) | 29.87 - 1.95<br>(2.02 - 1.95) |
| **Space group** | P 63 2 2 | P 41 21 2 |
| **Unit cell** | 106.8 106.8 136.9 | 69.4 69.4 150.6 |
| **Total reflections** | 115489 (4902) | 622246 (62705) |
| **Unique reflections** | 12644 (633) | 27623 (2366) |
| **Multiplicity** | 9.1 (7.7) | 22.5 (23.4) |
| **Completeness (%)** | 95.4 (76.9) | 94.95 (87.63) |
| **Mean I/sigma(I)** | 10.6 (1.4) | 6.28 (1.23) |
| **Wilson B-factor** | 24.50 | 19.50 |
| **R-merge** | 0.117 (1.579) | 0.2789 (2.671) |
| **R-meas** | 0.124 (1.692) | 0.2855 (2.73) |
| **R-pim** | 0.041 (0.595) | 0.06057 (0.5606) |
| **CC1/2** | 0.997 (0.563) | 0.993 (0.657) |
| **CC\*** | 1.000 (0.986) | 0.998 (0.891) |
| **Reflections used in refinement** | 12633 (68) | 26258 (2366) |
| **Reflections used for R-free** | 652 (3) | 1370 (115) |
| **R-work** | 0.1990 (0.3229) | 0.1718 (0.2928) |
| **R-free** | 0.2518 (0.3636) | 0.2032 (0.3389) |
| **CC(work)** | 0.949 (0.949) | 0.938 (0.754) |
| **CC(free)** | 0.923 (0.976) | 0.899 (0.699) |
| **Number of non-hydrogen atoms** | 2539 | 2698 |
| **Macromolecules** | 2393 | 2430 |
| **Ligands** | 0 | 0 |
| **Solvent** | 146 | 268 |
| **Protein residues** | 315 | 314 |
| **RMS(bonds)** | 0.002 | 0.014 |
| **RMS(angles)** | 0.45 | 1.08 |
| **Ramachandran favored (%)** | 97.09 | 98.37 |
| **Ramachandran allowed (%)** | 2.91 | 1.63 |
| **Ramachandran outliers (%)** | 0 | 0.00 |
| **Rotamer outliers (%)** | 2.35 | 1.12 |
| **Clashscore** | 4.18 | 3.10 |
| **Average B-factor** | 35.54 | 26.73 |
| **Macromolecules** | 35.93 | 25.80 |
| **Solvent** | 29.25 | 35.17 |

Statistics for the highest-resolution shell are shown in parentheses.

The structure revealed that the NirD domain adopts a classic Rossmann-like fold with six parallel beta sheets and sixteen alpha helices connected by loops **(Fig. 1B)**. Consistent with AlphaFold predictions, the annotated DDHD domain did not form a complete structural unit. Instead, the DDHD region constituted approximately one half of the NirD domain, with the remainder contributed by the preceding ∼200 amino acids **(Fig. 1C, 1D)**. Thus, the DDHD domain is not a bona fide standalone domain and will be referred to as the DDHD region. Notably, a similar architecture is predicted for other DDHD- containing proteins. For example, members of the intracellular phospholipase A1 (iPLA_1_) family are predicted to combine the DDHD region with adjacent sequences to form a complete domain **(Supplementary Fig. 1A-1C).**

### The NirD domain forms a dimer

The crystal structure further revealed the Nir2-NirD^LL^ domain assembles into a homodimer with a head-to-head configuration **(Fig. 2A)**. The dimer interface primarily involves hydrophobic interactions, with most intermolecular contacts formed by the α3 and α5 helices of one subunit packing against the α8 helix of the opposing subunit **(Fig. 2A, 2B)**. Additional interactions further stabilize the interface, creating an extensive contact surface between the two subunits. Notably, the N- and C-termini of each NirD monomer are positioned distal to the dimer interface and lie in close proximity to each other, separated by only ∼13.5 Å **(Fig. 2A)**. This structural arrangement indicates that the NirD domain contributes minimally to the overall length of full-length Nir2. Consequently, dimerization through the NirD domain is unlikely to substantially influence the distance that Nir2 can span between the ER and the PM at ER-PM junctions. The extensive dimer interface and compact organization of the termini suggest that the primary function of the NirD domain is to mediate protein-protein interactions rather than to serve as a structural spacer.

**Fig. 2.**
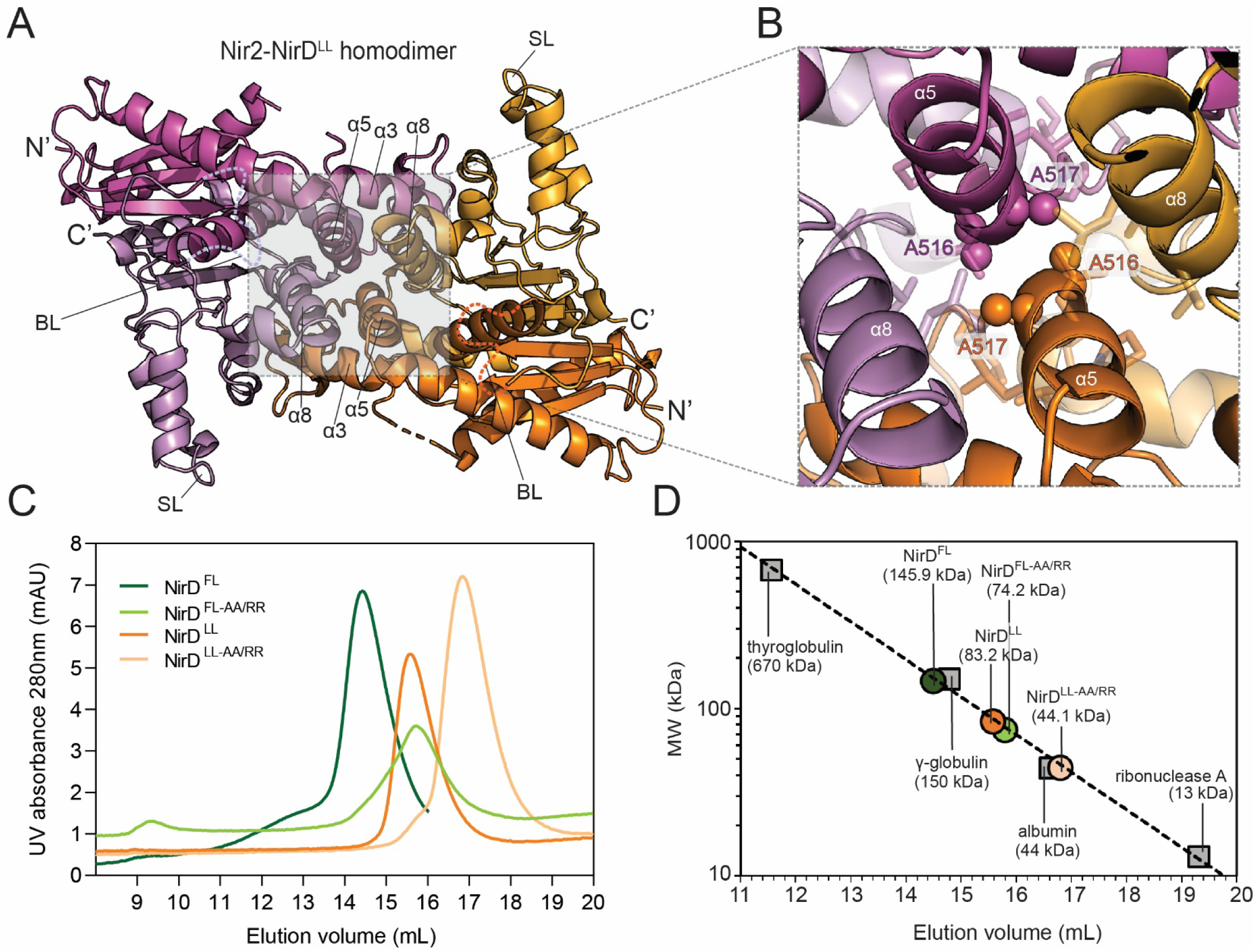
Arginine mutations at the dimer interface of Nir2 NirD domain produce a stable monomer. **(A)** Structure and orientation of the Nir2-NirD^LL^ homodimer. The dimer adopts a “head-to-head” orientation with the N- and C-termini located near each other and facing away from the dimer interface. Helices α3, α5, and α8 form the hydrophobic dimer interface. **(B)** Closer view of the homodimer interface and the interacting hydrophobic helices. Consecutive alanine residues on the α5 helix of each monomer lie directly transverse from one another and are candidates for mutation to dissociate the dimer *in vitro*. **(C)** Size exclusion chromatograms displaying the retention volumes of Nir2 NirD constructs before and after mutation. Substituting arginine residues in place of Ala516 and Ala517 (AA/RR) in the wild-type (WT) and loopless (LL) constructs increases their retention volumes, which indicates a decrease in apparent molecular weight (MW^app^). **(D)** Nir2 NirD MW^app^s interpolated from a standard curve derived from globular proteins of known MWs. The NirD^WT^ and NirD^LL^ MW^app^s decrease by half upon mutation.

### The Nir2 NirD domain does not bind membranes

Membrane association is important for Nir2 function. The C-terminal PA-binding LNS2 domain is both necessary and sufficient for PM recruitment of Nir2 ^6,19^, whereas the N-terminal PITP domain transiently interacts with membranes to mediate lipid transfer at ER-PM junctions. We therefore asked whether the NirD domain also contributes to membrane binding. To test this possibility, we generated liposomes with varying lipid compositions and assessed NirD membrane association using liposome co-sedimentation assays. Under all conditions examined, we observed minimal co-sedimentation of the Nir2 NirD domain with liposomes **(Supplementary Fig. 2)**, indicating little or no detectable membrane association. These results indicate that the Nir2 NirD domain does not directly bind membranes.

### NirD domains do not exhibit detectable esterase activity

Given the structural similarity between the NirD domain and the DDHD-family lipase/esterases **(Supplementary Fig. 1)**, we next tested if the Nir2 NirD domain possesses esterase activity. To assess catalytic activity, we employed the generic esterase substrate 4-nitrophenyl acetate (4-NPA) **(Supplementary Fig. 3A)**. As positive and negative controls, we used purified human DDHD2 and the catalytically inactive DDHD2 S351A mutant ^20,21^, in which the catalytic serine residue is replaced by alanine **(Supplementary Fig. 1D)**.

Under our assay conditions, the Nir2 NirD^LL^ domain exhibited no detectable esterase activity, comparable to the inactive DDHD2 S351A control **(Supplementary Fig. 3B)**. In contrast, wild-type DDHD2 efficiently hydrolyzed the ester bond of 4-NPA, confirming the validity of the assay. Similarly, the Nir1 NirD^LL^ domain displayed no detectable esterase activity **(Supplementary Fig. 3B)**. These results indicate that NirD domains lack intrinsic esterase activity. This conclusion is consistent with sequence and structural analyses showing that both Nir1 and Nir2 diverge from the catalytic machinery found in DDHD-family esterases. Notably, the catalytic serine residue is replaced by cysteine in Nir1 and glycine in Nir2, and the conserved histidine and aspartate residues that complete the catalytic triad are positioned too far apart to form the productive hydrogen-bonding network required for catalysis **(Supplementary Fig. 1D)**. Together, these observations support the conclusion that NirD domains are catalytically inactive despite their structural resemblance to DDHD-family enzymes.

### Point mutations at the dimer interface convert the Nir2 NirD domain to a monomer

Given the observation of a NirD homodimer in the crystal structure, we hypothesized that the primary function of the Nir2 NirD domain is to mediate homo- or heterodimerization among Nir family proteins. As an initial test of this hypothesis, we sought to determine whether the Nir2 NirD domain also exists as a dimer in solution.

To investigate this, we performed size-exclusion chromatography (SEC) to determine the oligomeric states of purified Nir2 NirD proteins. SEC does not quantitate absolute molecular weight (MW) but instead measures the hydrodynamic radius of a protein in solution, which can be correlated to an apparent MW (MW^app^) using a standard curve from the elution volumes of globular proteins with known MWs. The MW^app^ is sensitive to protein shape, and the presence of intrinsically disordered loops in a protein, such as the NirD^WT^ domains that retain the big and small loops, and will increase the hydrodynamic radii, and subsequently the MW^app^.

After performing our SEC analysis, the resulting retention volume of the Nir2 NirD^WT^ domain suggested that it formed a dimer in solution **(Fig. 2C, 2D, Table 2)**. NirD dimerization was also consistent with the elution volume of the NirD^LL^ domain that lacks the intrinsically disordered loops, which had an MW^app^ almost identical to the absolute MW of the NirD^LL^ dimer **(Fig. 2C, 2D, Table 2)**.

**Table 2.** Oligomerization states of WT and AA/RR Nir2 NirD domains calculated from size-exclusion elution volumes.

| **Protein** | **monomer MW (kDa)** | **dimer MW (kDa)** | **elution volume (mL)** | **MW<sup>app</sup> (kDa)** | **oligomeric state** |
| --- | --- | --- | --- | --- | --- |
| NirD<sup>WT</sup> | 49.5 | 99.0 | 14.5 | 152.4 | dimer |
| NirD<sup>WT-AA/RR</sup> | 49.7 | 99.4 | 15.8 | 77.8 | monomer |
| NirD<sup>LL</sup> | 36.3 | 72.6 | 15.6 | 86.3 | dimer |
| NirD<sup>LL-AA/RR</sup> | 36.5 | 73.0 | 16.8 | 46.4 | monomer |

Next, we sought to introduce point mutations to dissociate the Nir2 NirD dimers. Structural analysis identified an interface between the two dimer subunits involving hydrophobic alpha helices bundled together with two sequential alanine residues (Ala516 and Ala517) situated transverse from one another at the center of this interface **(Fig. 2B)**. We hypothesized that substitution of Ala516 and Ala517 to arginine residues would disrupt the homodimer by generating both repulsive charge interactions and steric clashes. We generated these point mutants in Nir2-NirD^WT^ and Nir2-NirD^LL^ and purified these proteins for SEC analysis.

Using SEC, the mutant proteins, denoted Nir2-NirD^WT-AA/RR^ and Nir2-NirD^LL-AA/RR^, had longer retention volumes with MW^app^s consistent with monomers **(Table 2)**. In addition, the MW^app^s of the AA/RR point mutants were approximately half of the MW^app^ of their respective wild-type counterparts, thus denoting a shift from a dimer to a monomer in solution **(Fig. 2C, 2D, Table 2)**. Taken together, these results demonstrate that the Nir2 NirD domain can form a dimer in solution, and that the AA/RR point mutations prevent dimerization and produce a stable NirD monomer.

### Structure of the Nir1 NirD domain reveals a near-identical homodimer

Prior studies found that Nir1 and Nir2 directly interact through a region located between the FFAT motif and LNS2 domain of Nir1, and deletion of this region abolished Nir1-Nir2 binding^13^. Our structural analysis revealed that this interaction region is part of the NirD domain, raising the possibility that NirD mediates heterodimerization between Nir1 and Nir2.

To investigate this possibility, we purified and structurally characterized the corresponding NirD domain from Nir1. As with the Nir2 NirD domain, we were unable to crystallize the complete Nir1 NirD domain and thus generated a “loopless” version denoted Nir1-NirD^LL^ **(Fig. 3A)**. Nir1-NirD^LL^ successfully crystallized, and the structure was refined to 1.95 Å **(Table 1).**

**Fig. 3.**
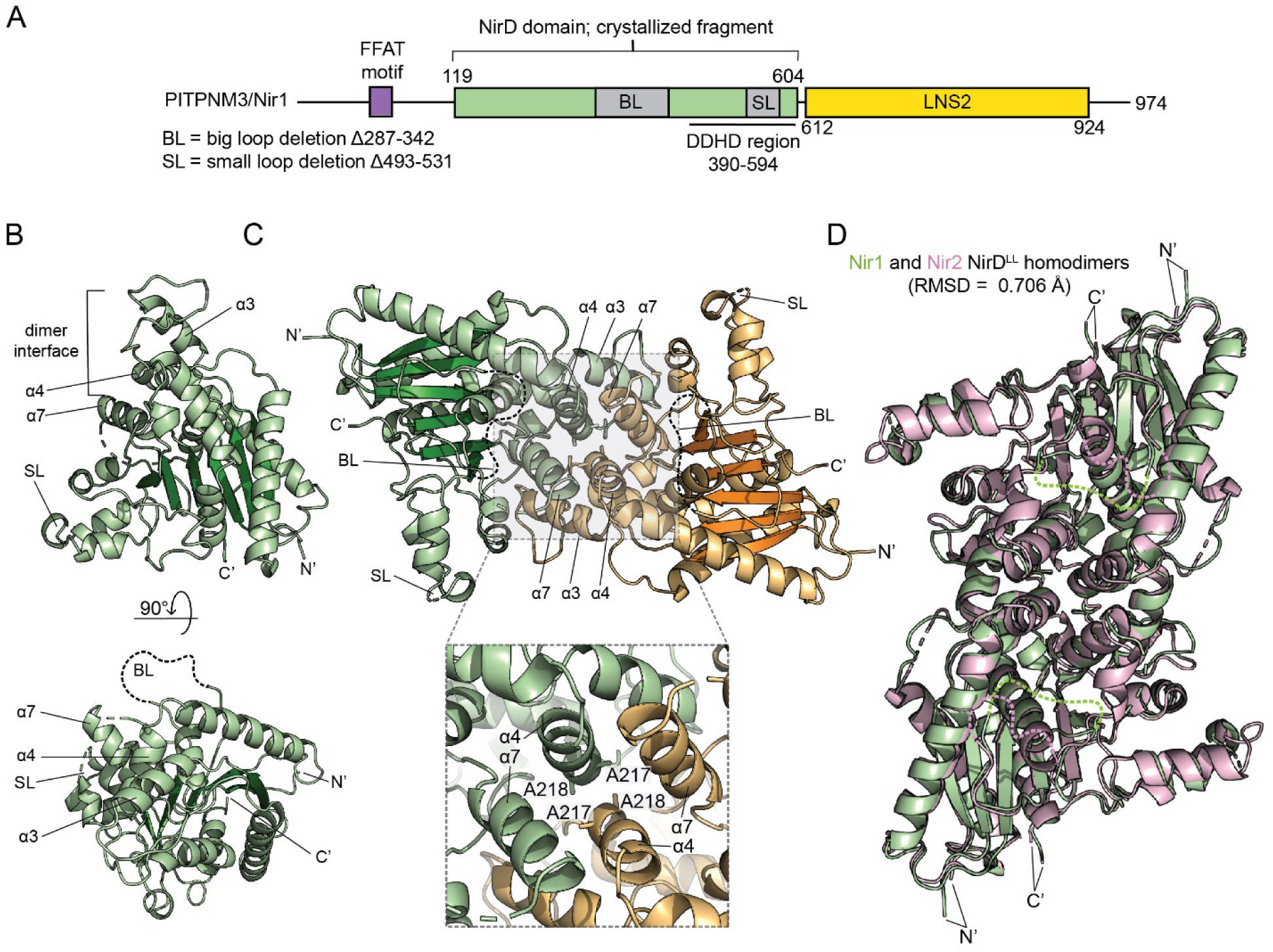
Crystal structure of the Nir1 NirD domain. **(A)** Domain architecture of Nir1 depicting the domain constraints of the novel domain and the crystallized fragment, the FFAT motif, and the LNS2 domain. Predicted disordered loops (grey) were removed prior to crystallization. **(B)** Overall structure of the Nir1-NirD^LL^ domain. Similar to Nir2-NirD^LL^, Nir1-NirD^LL^ contains two internal predicted disordered loops that were removed prior to crystallization (loop regions denoted “BL” and “SL”). Front view (top) shows the dimer interacting region. **(C)** Nir1-NirD^LL^ forms a homodimer. The monomers are aligned in a “head-to-head” arrangement that places the N- and C-termini facing away from the interface. The dimer relies on hydrophobic interactions between the α3, α4, and α7, and contains conserved consecutive alanine residues (Ala217 and Ala218) also found in the Nir2-NirD^LL^ dimer interface. **(D)** Superposition of the Nir1-NirD^LL^ (green) and Nir2-NirD^LL^ (pink) homodimers. The homodimers align with an RMSD value of 0.706 Å (PyMol).

The structure revealed that the Nir1-NirD^LL^ domain also forms a homodimer with each subunit being near-identical to the Nir2-NirD^LL^ structure (RMSD = 0.580 Å) **(Fig. 3B, 3C)**, indicating a high degree of structural conservation between Nir1 and Nir2 NirD domains. The overall dimer architecture is likewise highly conserved, as the orientation of the two subunits and the composition of the dimer interface in the Nir1 NirD^LL^-NirD^LL^ homodimer are nearly identical to those of Nir2 NirD^LL^-NirD^LL^ homodimer (RMSD = 0.706 Å) **(Fig. 3D)**. Notably, the pair of alanine residues located within the helical bundle at the dimer interface is conserved between Nir1 and Nir2 and adopts an identical orientation in both crystal structures **(Fig. 3C).** These observations suggest that the dimerization mechanism is conserved across the two paralogs and predict that the corresponding AA/RR substitutions would disrupt NirD-mediated dimerization in both Nir1 and Nir2.

### Full-length Nir1 and Nir2 proteins preferentially form heterodimers via the NirD domain in cells

Our crystal structures demonstrate that the NirD domains of Nir1 and Nir2 readily form homodimers. However, previous co-immunoprecipitation (co-IP) experiments showed that Nir1 preferentially interacts with Nir2 rather than self-associate ^13^, suggesting that Nir1-Nir2 heterodimers may be favored. Directly testing this *in vitro* is challenging because NirD homodimers must first dissociate into monomers before heterodimer formation can occur. To address this, we used a computational approach. AlphaFold2 Multimer predicted a Nir1-Nir2 heterodimer **(Fig. 4A)** that closely matches the Nir1 and Nir2 NirD homodimer crystal structures, with root mean square deviations (RMSDs) of 0.672 Å and 0.893 Å for the Cα backbone atoms. This strong structural similarity supports a shared interaction interface and suggests that the NirD domain can mediate Nir1-Nir2 heterodimerization. We therefore tested whether Nir1 and Nir2 preferentially form heterodimers in cells. All cellular experiments described below were performed using full-length human Nir1 and Nir2 proteins rather than the isolated NirD domains.

**Fig. 4.**
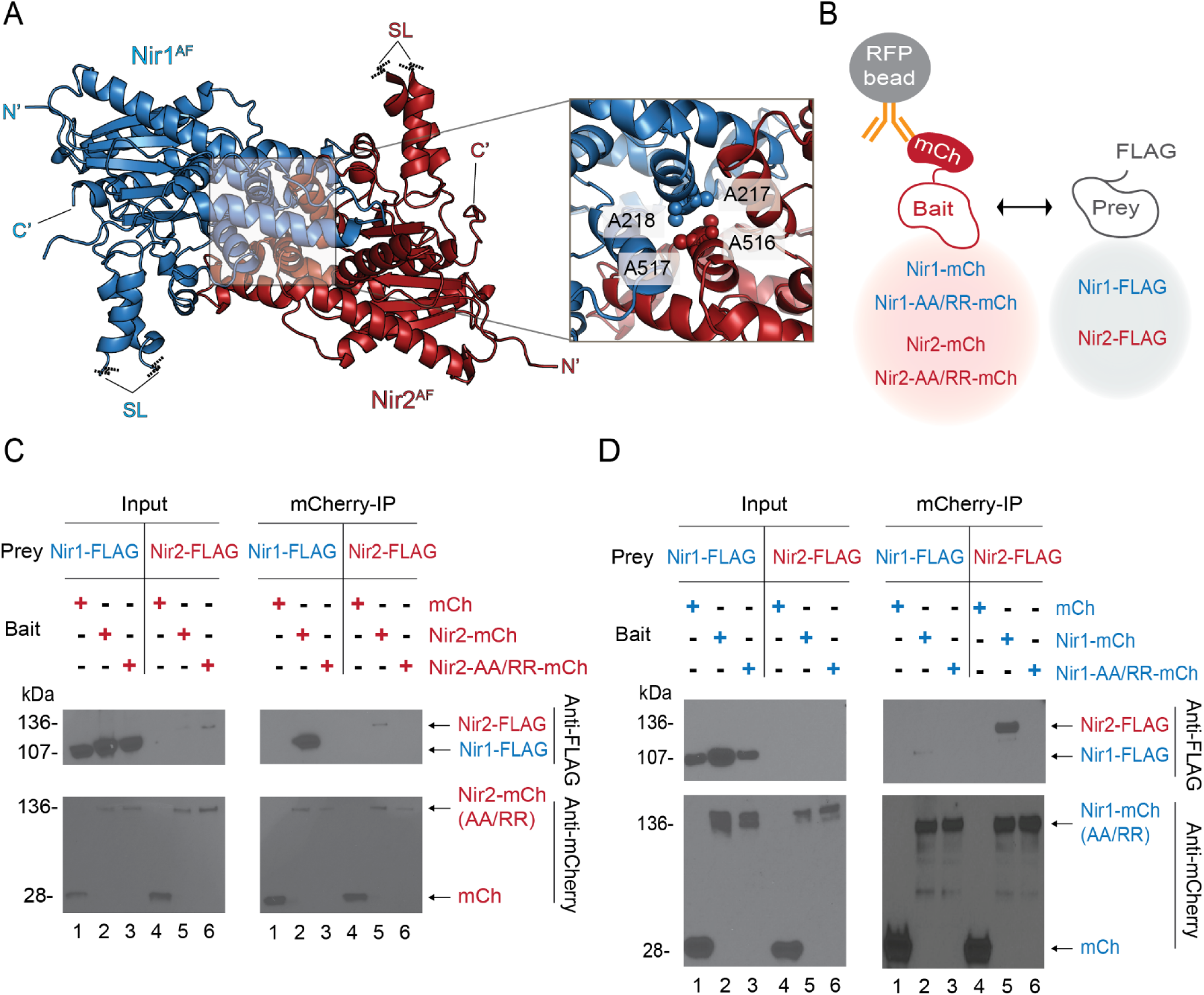
Preferential heterodimerization of Nir1 and Nir2 in cells is mediated by the NirD domain. **(A)** AlphaFold2 prediction of the Nir1-Nir2 heterodimer with the NirD domains in an identical orientation as the experimental Nir1 and Nir2 NirD homodimer crystal structures with the two Alanine residues of Nir1 (A217, A218) and Nir2 (A516, A517) at the central interface. The big loop (BL) and small loop (SL) are hidden for clarity. **(B)** Schematic of co-immunoprecipitation experiments used to assess the effects of point mutations on homo- and heterodimerization. **(C)** Co-immunoprecipitation of Nir1-FLAG or Nir2-FLAG with mCherry vector, Nir2-mCh, or Nir2-AA/RR-mCh co-expressed in HeLa cells. Representative blot from three independent experiments. Lane numbers are indicated below. **(D)** Co-immunoprecipitation of Nir1-FLAG or Nir2-FLAG with mCherry vector, Nir1-mCh, or Nir1-AA/RR-mCh co-expressed in HeLa cells. Representative blot from three independent experiments. Lane numbers are indicated below.

We performed co-IP experiments in HeLa cells co-transfected with mCherry-tagged Nir1 or Nir2 as “Bait” and FLAG-tagged Nir1 or Nir2 as “Prey” **(Fig. 4B)**. Consistent with our previous report ^13^, FLAG-tagged Nir1 (Nir1-FLAG) robustly co-immunoprecipitated with mCherry-tagged Nir2 (Nir2-mCh) (mCherry-IP lane #2, **Fig. 4C)**, but showed only weak association with Nir1-mCh (mCherry-IP lane #2, **Fig. 4D)**. Conversely, FLAG-tagged Nir2 (Nir2-FLAG) preferentially associated with Nir1-mCh over Nir2-mCh (mCherry-IP lane #5, **Fig. 4C, 4D)**, despite its low abundance in the input. These results demonstrate that, in cells and in the context of full-length proteins, Nir1 and Nir2 strongly favor heterodimer formation over homodimerization, even though their NirD domains can form homodimers *in vitro* at high micromolar concentrations.

To directly test whether the NirD domain mediates Nir1-Nir2 heterodimerization, we introduced AA/RR mutations into the NirD domains of mCherry-tagged Nir1 and Nir2. These mutations are designed to disrupt the dimer interface. As expected, Nir1-FLAG failed to co-immunoprecipitate with Nir2-AA/RR-mCh (mCherry-IP lane #3, **Fig. 4C**). Similarly, Nir2-FLAG did not co-immunoprecipitate with Nir1-AA/RR-mCh (mCherry-IP lane #6, **Fig. 4D)**. Together, these results demonstrate that Nir1 and Nir2 form heterodimers in cells through their NirD domains.

### Heterodimerization with Nir1 promotes Nir2 recruitment to ER-PM junctions in receptor-stimulated cells

Translocation of Nir2 from the cytosol to ER-PM junctions is essential for Nir2 to drive the PI cycle in receptor-stimulated cells ^6,7^. We and others have shown that Nir2 recruitment to ER-PM junctions occurs only under strong stimulation conditions, such as receptor overexpression or exposure to high concentrations of PA ^6–8^. Importantly, we recently demonstrated that this requirement is bypassed by co-expression of Nir1 ^13^, indicating that Nir1 actively promotes Nir2 recruitment under physiological stimulation conditions.

To determine whether the NirD domain mediates Nir1-dependent Nir2 recruitment to ER-PM junctions, we examined the near-adhesion surface of HeLa cells co-transfected with Nir1-YFP and either Nir2-mCh or the dimerization-defective mutant Nir2-AA/RR-mCh using confocal microscopy. In contrast to wild-type Nir2 **(Fig. 5A)**, the Nir2-AA/RR mutant showed markedly reduced translocation to ER-PM junctions following histamine stimulation **(Fig. 5B, 5D).** Consistently, co-expression of Nir2-mCh with the Nir1-AA/RR mutant resulted in a predominantly cytosolic distribution of Nir2 and significantly impaired recruitment to ER-PM junctions compared with cells expressing wild-type Nir1 **(Fig. 5C, 5D)**.

**Fig. 5.**
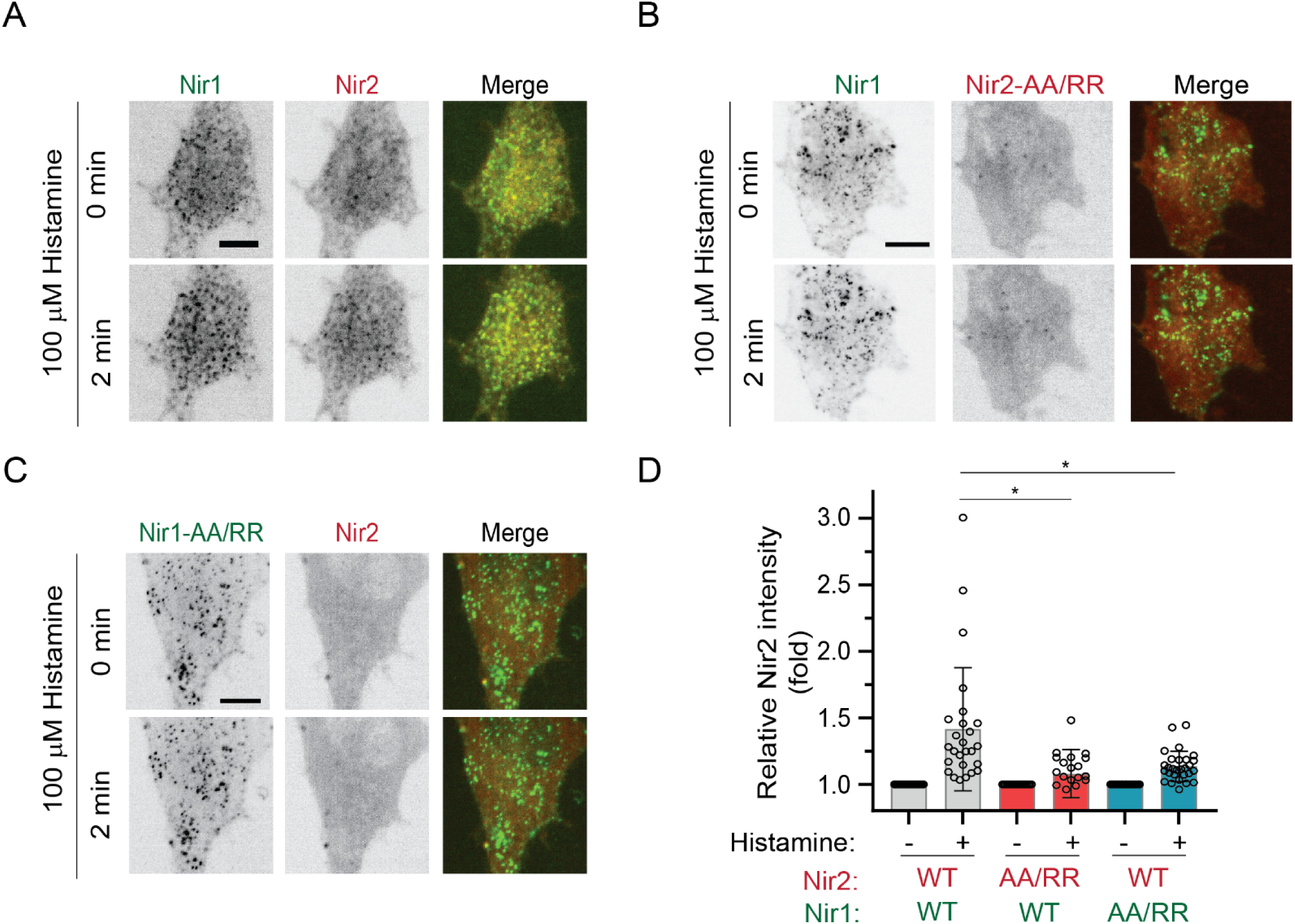
Nir1-driven recruitment of Nir2 to ER-PM junctions in receptor-stimulated cells requires Nir1-Nir2 heterodimerization via the NirD domain. **(A)** Confocal images of HeLa cells co-expressing Nir1-YFP and Nir2-mCh following stimulation with 100 μM histamine for the indicated times. Scale bar, 10 µm. **(B)** Confocal images of HeLa cells co-expressing Nir1-YFP and Nir2-AA/RR-mCh under the same conditions as in (A). Scale bar, 10 µm. **(C)** Confocal images of HeLa cells co-expressing Nir2-YFP and Nir1-AA/RR-mCh under the same conditions as in (A). Scale bar, 10 µm **(D)** Quantification of Nir2 recruitment to ER-PM junctions as relative changes in fluorescence intensity of Nir2-mCh, Nir2-AA/RR-mCh, and Nir2-YFP in HeLa cells co-expressing Nir1-YFP or Nir1-AA/RR-mCh following histamine stimulation. Data represent mean ± SEM (* *p* < 0.05, t test; Nir2-WT n = 26; Nir2-AA/RR n = 21; Nir1-AA/RR n = 28; three independent experiments).

Importantly, the AA/RR mutations did not disrupt Nir1 pre-localization to ER-PM junctions **(Supplementary Fig. 4A)**, or Nir2 recruitment under strong stimulation conditions in receptor-overexpressing cells **(Supplementary Fig. 4B)**. Thus, these mutations specifically impair Nir1-dependent recruitment of Nir2 without affecting the intrinsic ability of either protein to localize at ER-PM junctions. Together, these results demonstrate that Nir1-driven recruitment of Nir2 to ER-PM junctions in receptor-stimulated cells requires Nir1-Nir2 heterodimerization mediated by the NirD domain.

### Nir1-Nir2 heterodimerization facilitates PIP_2_ replenishment during the PI cycle

We next directly examined the functional importance of NirD-mediated Nir1-Nir2 heterodimerization in the PI cycle. We previously developed a total internal reflection fluorescence (TIRF) microscopy assay to monitor the consumption and replenishment of PM PIP_2_ during the PI cycle in living HeLa cells ^7,23^. In this assay, cells are co-transfected with the histamine H1 receptor (H1R) to enhance PLC-mediated PIP_2_ hydrolysis and the PIP_2_ biosensor GFP-PLCδ-PH to track changes in PM PIP_2_ levels in real time. Using this approach, we previously demonstrated that knockdown of either Nir1 or Nir2 markedly impairs PIP_2_ replenishment in receptor-stimulated cells ^7,13^. Based on these observations, we hypothesized that Nir1 and Nir2 cooperatively promote the PI cycle through NirD-mediated heterodimerization.

To test this hypothesis, we modified the assay to enable simultaneous expression of four components (Nir1, Nir2, H1R and GFP-PLCδ-PH) in the same cells. HeLa cells were co-transfected with an internal ribosome entry site (IRES) construct that co-expresses H1R and GFP-PLCδ-PH at a fixed ratio, together with Nir1 and/or Nir2. Cells were stimulated with 10 μM histamine to induce robust PIP_2_ consumption followed by replenishment, which was monitored as changes in GFP-PLCδ-PH fluorescence using TIRF microscopy **(Fig. 6A)**. Overexpression of either Nir1 or Nir2 alone substantially enhanced PIP_2_ replenishment compared with the vector-transfected control **(Fig. 6B)**. Notably, co-expression of Nir1 and Nir2 produced a significantly greater effect than either protein alone, demonstrating that the two proteins act cooperatively to promote PIP_2_ replenishment during receptor signaling.

**Fig. 6.**
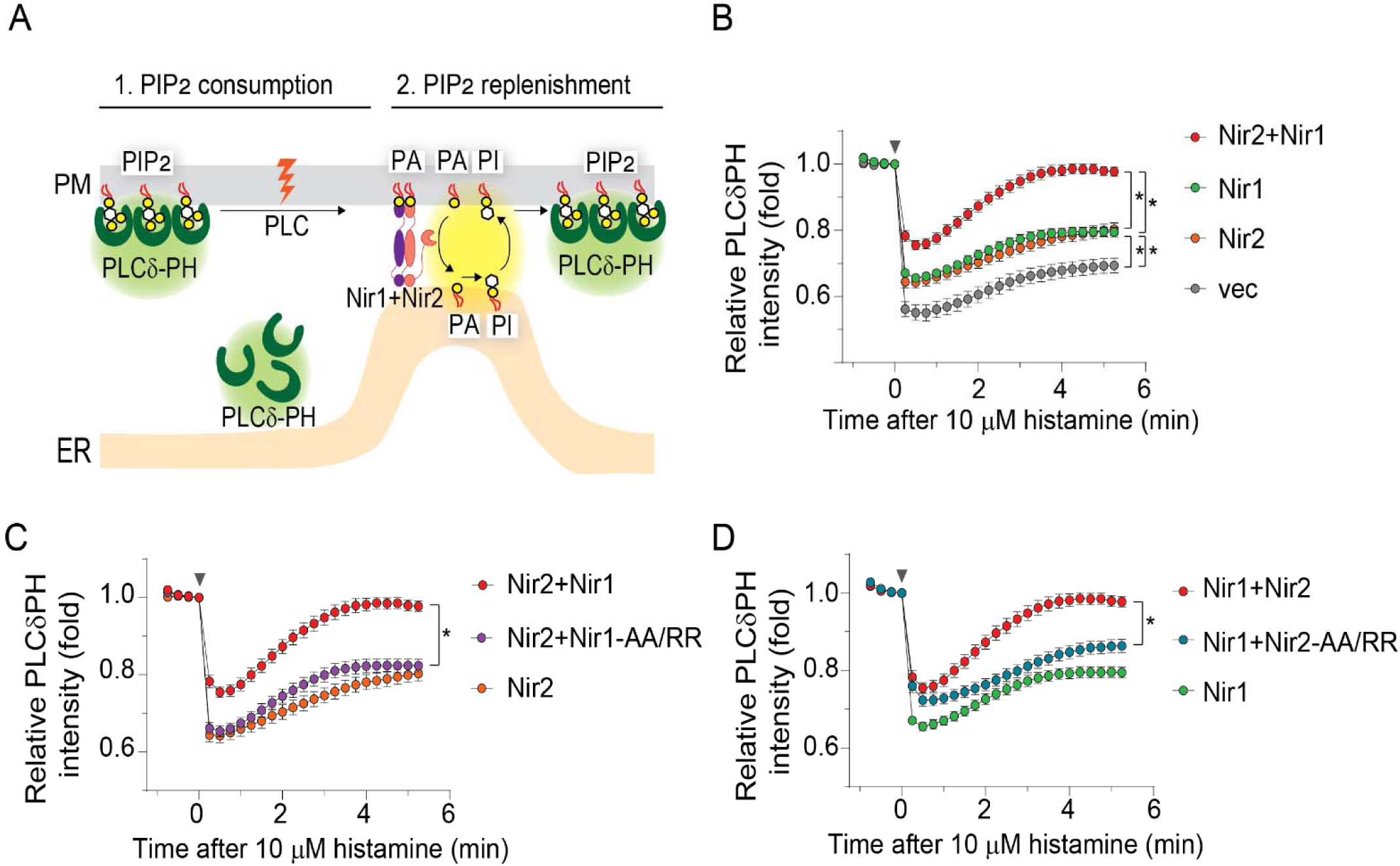
Nir1-Nir2 heterodimerization strongly enhances PIP_2_ replenishment in the PI cycle. **(A)** Schematic illustrating plasma membrane (PM) PIP_2_ consumption by PLC and subsequent replenishment by Nir1 and Nir2 in receptor-stimulated cells, monitored using the PIP_2_ biosensor GFP-PLCδ-PH. **(B)** Time course of PM PIP_2_ changes following stimulation with 10 μM histamine in HeLa cells co-transfected with H1R-IRES-GFP-PLCδ-PH and either Nir2-mCherry + Nir1-mTagBFP2, Nir1-mTagBFP2 alone, Nir2-mCherry alone, or vector control, measured by by TIRF microscopy. Arrowhead indicates histamine addition. Data represent mean ± SEM (> 20 cells from three independent experiments); * p < 0.05 (t test). **(C)** PM PIP_2_ dynamics following 10 μM histamine stimulation in cells co-expressing H1R-IRES-GFP-PLCδ-PH with Nir1-mTagBFP2 and either Nir2-mCherry, Nir2-AA/RR-mCherry, or vector control. Data represent mean ± SEM (> 30 cells from three independent experiments); * p < 0.05 between Nir1 + Nir2 and Nir1 + Nir2-AA/RR (t test). **(D)** PM PIP_2_ dynamics following 10 μM histamine stimulation in cells co-expressing H1R-IRES-GFP-PLCδ-PH with Nir2-mCherry and either Nir1-mTagBFP2, Nir1-AA/RR-mTagBFP2, or vector control. Data represent mean ± SEM (> 20 cells from three independent experiments); * p < 0.05 between Nir2 + Nir1 and Nir2 + Nir1-AA/RR (t test).

To determine whether this cooperative effect depends on NirD-mediated heterodimerization, we repeated these experiments using the AA/RR dimerization-defective mutants. In contrast to wild-type Nir1, the Nir1-AA/RR mutant failed to further enhance PIP_2_ replenishment when co-expressed with Nir2 **(Fig. 6C)**. Similarly, disruption of the NirD dimerization interface in Nir2 significantly reduced its ability to potentiate PIP_2_ replenishment when co-expressed with Nir1 **(Fig. 6D)**. Together, these findings establish NirD-mediated heterodimerization as the molecular basis of Nir1-Nir2 cooperation and demonstrate that it is a key mechanism for enhancing PI cycle activity and maintaining phosphoinositide homeostasis during receptor signaling.

### Dimerization with Nir1 expands the sensitivity and dynamic range of Nir2 recruitment to ER-PM junctions

The PI cycle sustains cellular signaling and homeostasis in response to a wide range of PLC-activating stimuli, including histamine, growth factors, antigens, and neurotrophins ^3–5^. To function effectively under these diverse signaling conditions, the PI cycle must be both sensitive to weak inputs and capable of scaling its output in response to stronger stimulation. We hypothesized that dimerization with Nir1 enables Nir2 to operate across a broad range of stimulation intensities, thereby enhancing the robustness of the PI cycle. This hypothesis is supported by previously findings that the LNS2 domain of Nir1 exhibits greater sensitivity to PA than the corresponding domain of Nir2 ^24^, and that Nir1 displays stronger localization to ER-PM junctions than a comparable Nir2 construct lacking the PITP domain (Nir2-ΔPITP) ^13^.

To test this hypothesis, we quantified the recruitment of Nir2 and the dimerization-defective Nir2-AA/RR mutant to ER-PM junctions in HeLa cells co-expressing Nir1 and H1R. Cells were stimulated with histamine concentrations spanning five orders of magnitude (100 pM to 10 μM), and Nir2 protein recruitment was monitored by TIRF microscopy, which provides a sensitive measure of protein accumulation at ER-PM junctions ^23^. Changes in fluorescence intensity were quantified 1 minute after stimulation. In cells co-expressing Nir1 and H1R, robust recruitment of Nir2 was detected at histamine concentrations as low as 1 nM **(Fig. 7A, 7B).** Moreover, increasing histamine concentrations produced a progressive increase in Nir2 recruitment across the entire concentration range tested, without evidence of saturation **(Fig. 7B)**. These results demonstrate that Nir2 exhibits both high sensitivity and a broad dynamic range when partnered with Nir1. In contrast, the Nir2-AA/RR mutant, which is unable to dimerize with Nir1, showed detectable recruitment only at histamine concentrations of 100 nM or greater **(Fig. 7C, 7D).** Furthermore, recruitment of Nir2-AA/RR reached a plateau at high histamine concentrations (1-10 μM), indicating a substantially restricted dynamic range. Together, these findings demonstrate that Nir1-Nir2 heterodimerization expands both the sensitivity and dynamic range of Nir2 recruitment to ER-PM junctions. By lowering the threshold for Nir2 recruitment while simultaneously increasing its responsiveness to progressively stronger stimulation, Nir1 enables graded accumulation of Nir2 in proportion to signaling demand. This mechanism provides a direct molecular explanation for how the PI cycle maintains efficient phosphoinositide replenishment across of a broad spectrum of stimulus intensities.

**Fig. 7.**
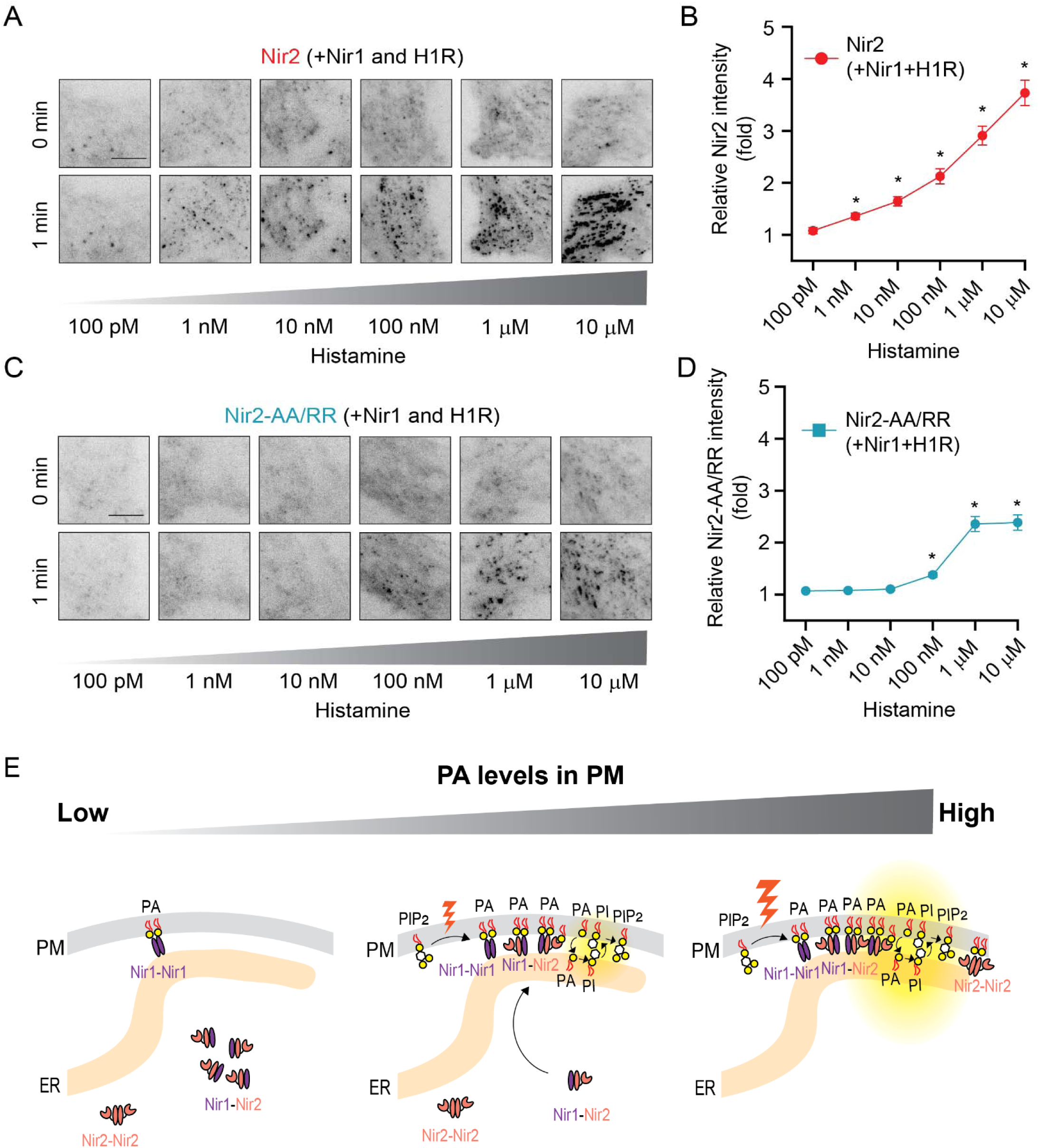
Dimerization with Nir1 expands the sensitivity and dynamic range of Nir2 recruitment to ER-PM junctions in receptor-stimulated cells. **(A)** TIRF images of HeLa cells co-expressing H1R, Nir1-YFP, and Nir2-mCh before and after stimulation with the indicated concentrations of histamine. Scale bar, 10 µm. **(B)** Quantification of Nir2-mCh recruitment to ER-PM junctions, shown as relative changes in fluorescence intensity in regions marked by Nir1-YFP, following histamine stimulation as in (A). Data represent mean ± SEM (40–50 cells per condition from three independent experiments); *p < 0.05 compared with 100 pM (t test). **(C)** TIRF images of HeLa cells co-expressing H1R, Nir1-YFP, and Nir2-AA/RR-mCh before and after stimulation with the indicated concentrations of histamine. Scale bar, 10 µm. **(D)** Quantification of Nir2-AA/RR-mCh recruitment to ER-PM junctions under the same conditions as in (C). Data represent mean ± SEM (40–50 cells per condition from three independent experiments); *p < 0.05 compared with 100 pM (t test). **(E)** Model of Nir1-Nir2 heterodimerization conferring robustness to the PI cycle. See text.

Taken together, our findings support a model in which Nir1-Nir2 heterodimerization confers robustness to the PI cycle **(Fig. 7E)**. Structural and biochemical analyses indicate that Nir1 and Nir2 preferentially form heterodimers through the NirD domain, favoring the formation of Nir1-Nir2 complexes over homodimers in cells. Under resting conditions, low PM PA levels preferentially support the localization of Nir1-Nir1 homodimers at ER-PM junctions owing to the higher PA sensitivity of Nir1. Following receptor stimulation, increased PA production drives proportional recruitment of Nir1-Nir2 heterodimers to ER-PM junctions, where Nir2 exchanges PA and PI to replenish PM PIP_2_. Through this mechanism, Nir1 functions as a sensitizing and tuning factor that enables PI cycle activity to scale with signaling demand. In the absence of Nir1-mediated heterodimerization, Nir2 recruitment is restricted to strong stimulation conditions, resulting in reduced sensitivity and a narrower operation range of the PI cycle.

## DISCUSSION

In this study, we uncover a fundamental mechanism by which Nir1 regulates Nir2 to confer robustness in the PI cycle. We identify a conserved Nir dimerization (NirD) domain that drives preferential Nir1-Nir2 heterodimerization in cells. In addition, we demonstrate that NirD-mediated Nir1-Nir2 dimerization is required for efficient recruitment of Nir2 to ER-PM junctions and promotes PM PIP_2_ replenishment in receptor-stimulated cells. Importantly, Nir1-Nir2 heterodimerization expands both the sensitivity and dynamic range of Nir2 recruitment, enabling graded and scalable responses across a broad spectrum of PLC-activating inputs that sustain cellular signaling and homeostasis ^3–5^. These findings reveal the structural basis of Nir1-dependent Nir2 recruitment and establish paralog heterodimerization as a key regulatory mechanism underlying PI cycle function.

A major advance of this work is the identification of the NirD domain and the consequent reinterpretation of the previous annotated DDHD domain. Our structural and biochemical analyses show that the DDHD region, previously of unclear function, has been repurposed from an enzymatic fold into a protein-protein interaction module. Although homologous to the catalytic domains of DDHD1 and DDHD2 lipases ^20,21,25,26^, the Nir DDHD region lacks enzymatic activity and instead cooperates with adjacent sequences to form the NirD dimerization domain. This functional transition illustrates how catalytic scaffolds can be adapted for regulatory roles at membrane contact sites.

Our structural analyses further reveal that the NirD domain forms highly conserved homodimers in both Nir1 and Nir2, whereas cellular studies demonstrate that full-length Nir1 and Nir2 preferentially assemble into heterodimers. These findings provide a molecular explanation for previous observations that a region between the FFAT motif and LNS2 domain is required for Nir1 interaction with Nir2 ^13^. More broadly, they support a model in which the two paralogs have functionally specialized following gene duplication: Nir2 retains lipid transfer activity, whereas Nir1 acts as a non-catalytic regulator that controls Nir2 localization and recruitment. Heterodimerization therefore couples localization to function, creating a complex that can efficiently and rapidly mobilize PI/PA exchange activity to ER-PM junctions in response to receptor stimulation.

Our findings also highlight distinct regulatory strategies within the Class IIA PITP family. Previous studies suggest that Nir3 functions independently of Nir2 and operates under low-stimulation conditions in a tissue-restricted manner ^6,12^. In contrast, Nir2 functions cooperatively with Nir1 to support PI cycle activity across a broad range of signaling conditions. These observations reveal functional specialization within the Nir protein family and underscore the importance of regulatory diversification in maintaining phosphoinositide homeostasis.

A key conceptual advance of this work is the demonstration that Nir1 enables stimulus-strength-dependent recruitment of Nir2. In the presence of Nir1, Nir2 is recruited efficiently at low levels of stimulation and continues to accumulate over a broad range of stimulus intensities without reaching saturation. In contrast, disruption of heterodimerization markedly increases the stimulation threshold required for Nir2 recruitment and restricts its maximal response. Therefore, Nir1 does not simply increase the amount of Nir2 recruitment to ER-PM junctions. Instead, it enables graded recruitment of Nir2 in proportion to signaling demand, thereby expanding both the sensitivity and dynamic range of the PI cycle.

This mechanism addresses a central challenge in lipid signaling: maintaining PIP_2_ homeostasis despite rapid and highly variable rates of consumption ^3–5^. PLC activation can vary substantially depending on receptor identity, ligand concentration, and cellular context, creating corresponding demands for phosphoinositide replenishment. By lowering the activation threshold for Nir2 recruitment while preserving responsiveness at higher stimulation levels, Nir1 effectively functions as a gain-control mechanism that matches PI transport capacity to signaling demands. In this way, Nir1-Nir2 heterodimerization provides a direct molecular basis for the robustness of the PI cycle, ensuring efficient PIP_2_ replenishment across a broad range of physiological conditions. In the absence of Nir1-mediated regulation, PI cycle activity is restricted to strong stimulation conditions, narrowing the range over which cells can maintain signaling competence.

Our findings may also provide mechanistic insights into Nir1-associated human diseases. Mutations in Nir1 have been linked to inherited retinal dystrophy ^14,16,17^, whereas elevated Nir1 expression has been associated with cancer progression and metastasis ^27–29^. Because Nir1 lacks lipid transfer activity, the molecular basis of these phenotypes has remained difficult to reconcile with the established role of Nir2 as the principal lipid transfer protein of the PI cycle. Our results suggest that disruption of Nir1 function may impair phosphoinositide homeostasis indirectly by altering the recruitment dynamics of Nir2 and limiting the ability of cells to adapt to changing signaling demands. Future studies examining whether disease-associated mutations affect NirD structure, dimerization, or stimulus-dependent Nir2 recruitment may provide important insights into disease pathogenesis.

More broadly, our study highlights the importance of protein-protein interactions at ER-PM junctions in regulating lipid homeostasis. Membrane contact sites are increasingly recognized as dynamic signaling platforms whose activities must be continuously adjusted to changing physiological demands. The Nir1-Nir2 complex exemplifies how scaffolding interactions can tune both the timing and magnitude of lipid transfer to match cellular requirements. Similar dimerization mechanisms may operate in other membrane contact site systems ^30^, representing a general strategy for coordinating lipid metabolism with cellular signaling.

In summary, we identify the NirD domain as a conserved dimerization module that mediates preferential Nir1-Nir2 heterodimerization and underlies Nir1-dependent recruitment of Nir2 to ER-PM junctions. By promoting efficient PIP_2_ replenishment and enabling stimulus-strength-dependent recruitment of Nir2, Nir1-Nir2 heterodimerization expands both the sensitivity and dynamic range of PI cycle responses. These findings uncover a previously unrecognized mechanism for tuning lipid transfer capacity to signaling demands and establish paralog heterodimerization as a fundamental strategy for achieving robust phosphoinositide homeostasis.

## METHODS

### Recombinant protein plasmids

The full-length Nir2 gene (accession code: NP_001124320) was gifted from Dr. Tamas Balla and the full-length Nir1 gene (accession code: NC_000017.11) was codon optimized for *E. coli* expression and gene synthesized (Twist Bioscience) in the pET28a plasmid. DNA oligo primers were synthesized (Integrated DNA Technologies) for Nir1-NirD^WT^ using residues 120-605 and Nir2-NirD^WT^ using residues 420-887 and inserted into ppSUMO plasmids containing N-terminal His-tags, followed by a ULP1 cleavable SUMO protein. Point mutations and internal loop deletions were made in the ppSUMO plasmids using the Q5 Site-Directed Mutagenesis Kit (New England Biolabs; E0554). Disordered loop regions (residues 586-679 and 785-818) were deleted from Nir2-NirD^WT^ and residues 286-343 and 492-532 were deleted from Nir1-NirD^WT^ to generate the “loopless” (NirD^LL^) constructs. The full-length DDHD2 (accession code: NP_001157704.1) was codon optimized for expression in E. coli (Twist Biosciences) and gene synthesized in the pET28a plasmid. Point mutagenesis to generate the inactive mutant DDHD2-S315A plasmid used the Q5 Site-Directed Mutagenesis Kit described above. All constructs were verified with direct sequencing.

### NirD domain protein overexpression and purification

All NirD Nir1 and Nir2 plasmids were transformed into competent BL21 RIPL cells (Agilent Technologies, Cat. No. 230280) for protein overexpression. Cells were grown at 37°C to an OD_600_ of 1.5 and induced with 100 μM isopropyl β-D-1-thiogalactopyranoside (IPTG) at 15°C for overnight growth. Cell pellets were harvested and lysed via sonication in buffer comprised of 50 mM Tris pH 7.5, 500 mM NaCl, 5% glycerol, 1% Triton X-100, and 2 mM beta-mercaptoethanol (βME), and lysates were centrifuged at 82,000 xg at 4°C for 1 hour. Protein-rich supernatant was collected, and the affinity-tagged proteins were isolated using Ni-NTA gravity flow affinity chromatography and eluted with buffer comprised of 50 mM Tris pH 7.5, 500 mM NaCl, 300 mM imidazole pH 7.5, and 5 mM βME. Eluates were incubated overnight at 4°C with ULP1 protease to remove the SUMO fragment. Isolated proteins were applied to Superdex 75 26/60 HiLoad column (GE Healthcare) or Superdex 200 26/600 HiLoad column (Cytiva) equilibrated with buffer comprised of 20 mM Tris pH 7.5, 150 mM NaCl, 10 mM βME, and 1 mM dithiothreitol (DTT). Purified proteins were concentrated to 1-12 mg/mL, flash frozen in liquid nitrogen in 30 μL aliquots, and stored at −80 °C.

### DDHD2 protein overexpression and purification

Both WT and S351A DDHD2 plasmids were transformed into competent BL21 RIPL cells (Agilent Technologies, Cat. No. 230280) for protein overexpression. Cells were grown at 37°C to an OD_600_ of 1.5 and induced with 0.5 mM isopropyl β-D-1-thiogalactopyranoside (IPTG) at 15°C for overnight growth. Cell pellets were harvested and lysed by sonication in a buffer comprised of 50 mM Tris-HCl pH 8, 500 mM NaCl, 5% glycerol, 1 mM Tris(2-carboxyethyl)phosphine (TCEP), 1 mM PMSF. Lysates were centrifuged at 68000 xg at 4°C for 1 hr. Supernatant was collected and the His-tagged proteins were isolated using Ni-NTA chromatography and eluted in buffer comprised of 50 mM Tris-HCl pH 8, 500 mM NaCl, 5% glycerol, 1 mM TCEP, and 300 mM imidazole. Isolated DDHD2 proteins were applied to the Superdex 200 column equilibrated with buffer comprised of 20 mM Tris-HCl pH 8, 150 mM NaCl, 1 mM TCEP. The purified proteins were concentrated to 1-2 mg/mL, flash frozen in liquid nitrogen, and stored at −80 °C.

### Liposome co-sedimentation

Palmitoyl oleoyl (PO) phospholipids (Avanti Polar Lipids) dissolved in chloroform:methanol solution were dried under nitrogen gas and resuspended in buffer A comprised of 150 mM NaCl and 20 mM Tris pH 7.5 to generate a 2 mM solubilized lipid mixture. Solubilized PO lipids underwent five freeze/thaw cycles with liquid nitrogen and were then sonicated for two minutes in a water bath to generate liposomes. 50 μL of pure Nir2-NirD^WT^ at 0.8 mg/mL was incubated with 50 μL of liposome mixture (1 mM total lipids) for 30 minutes at room temperature before centrifuging reactions in a vacuum for 1 hr at 100,000 xg at 4°C. 75 μL of the supernatant (S) was collected for SDS-PAGE analysis, after which the remaining supernatant was discarded. The remaining liposome pellet (P) was resuspended in 100 μL of buffer A, and 75 μL was collected for samples to be resolved via SDS-PAGE. ImageJ software^31^ was used to quantify pixel intensity of the S and P fraction gel bands for each condition. % protein bound was determined using [P/(P+S)]*100.

### Esterase activity assay

Esterase activity was measured by the hydrolysis rate of a generic esterase substrate, 4-nitrophenyl acetate (4-NPA) (Sigma, catalog #M8130). When hydrolyzed by an enzyme with esterase activity, p-nitrophenol is released into solution, which is read by absorbance at 405 nm. A stock of 4-NPA (35 mM) was freshly prepared daily by dissolving 63 mg in 10 mL of methanol and was slowly added to the activity buffer to prevent precipitation to a final concentration of 250 μM. The activity assay buffer used was comprised of 50 mM HEPES pH 8, 150 mM NaCl, and 0.32% Triton X-100. Purified aliquots of the DDHD2 protein were used as a positive control to determine the linearity of the reaction over time, which was linear between 15 minutes and 60 minutes of total reaction time. The protein concentration linearity of DDHD2 was also determined, ranging from 0.1 μM to 1.2 μM. DDHD2-S351A, an inactive version of the enzyme^20,21^, was used as a negative control. Purified aliquots of DDHD2, DDHD2-S351A, Nir1-NirD^LL^ and Nir2-NirD^LL^ were analyzed at a concentration of 1 μM in a 100 μL reaction. The amount of p-nitrophenol produced in each condition was quantified using a standard curve of p-nitrophenol (Sigma, catalog #241326) determined using the activity assay buffer.

### Crystallization and data collection

Crystals of Nir2-NirD^LL^ (10 mg/mL) were grown by hanging drop vapor diffusion using a 1:1 ratio of protein to well solution, which was comprised of 0.2 M Li_2_SO_4_ and 8% (w/v) polyethylene glycol (PEG) 3350. A solution of 30% glycerol and 30% (w/v) PEG 3350 was used as a cryoprotectant. Crystals of Nir1-NirD^LL^ (10 mg/mL) were grown by hanging drop vapor diffusion using a 1:1 ratio of protein to well solution, which was comprised of 0.1 M Tris pH 7.5, 8.5% (v/v) isopropanol, 21% (v/v) glycerol, and 12% (w/v) PEG 4000. The same cryoprotectant solution as Nir2-NirD^LL^ was used. Native diffraction data were collected at the Brookhaven National Lab NSLS-II FMX beamline 17-ID-2. Diffraction data for Nir2-NirD^LL^ was processed using the autoPROC pipeline ^32^ with STARANISO ^33^. Diffraction data for Nir1-NirD^LL^ was processed using DIALS ^34^ and scaled using AIMLESS ^35^ in CCP4 ^36^.

### Structure determination and refinement

Phases were determined by molecular replacement in Phenix ^37^ using Phaser ^38^. A truncated Nir2-NirD^LL^ AlphaFold2 model with B-factor adjustments using Phenix was used as a search model. An initial model was generated using Phenix Autobuild ^39^. Additional model building in Coot ^40^ and refinements using Phenix Refine ^41^ produced the final model of Nir2-NirD^LL^ at 2.45 Å resolution **(Table 1, PDB code: 9C37)**. Residues 581-585 and 680-683 that link the deleted big loop region, and residues at both the N- and C-terminus, residues 420-424 and 880-887, were omitted from the final model due to a lack of electron density. Molecular replacement for Nir1-NirD^LL^ used the Nir2-NirD^LL^ structure as a search model. Additional model building in Coot ^40^ and refinement in Phenix ^37^ produced the final model **(Table 1, PDB code: 9C38)**. Residues 280-285 and 344-386 that link the deleted big loop region, residues 489-491 and 533-536 that link the deleted small loop region, and residues 416-423 were omitted from the final model due to a lack of electron density.

### SEC oligomerization determination

A Superdex 200 Increase 10/300 column (Cytiva) was calibrated using a protein standard mix 15-600 kDa (69385; Sigma Aldrich). 100 μg each of the Nir2 NirD constructs, NirD^WT^, NirD^WT-AA/RR^, NirD^LL^, and NirD^LL-AA/RR^, were applied to the column consecutively on the same day using identical buffers consisting of 20 mM Tris pH 7.5, 150 mM NaCl, 10 mM βME, and 10 mM DTT. The peak retention volumes of each construct were plotted against the standard calibration curve to determine their approximate molecular weight (MW^app^) and oligomerization state.

### Cell culture, transfection, and stimulation

HeLa cells (American Type Culture Collection) were maintained in Dulbecco’s Modified Eagle Medium (DMEM; high glucose, HyClone) supplemented with 10% fetal bovine serum (FBS; HyClone) and 1% penicillin–streptomycin at 37°C in a humidified incubator with 5% CO_2_. Plasmid DNA was transfected using TransIT-LT1 reagent (Mirus Bio) for 16–20 h prior to experiments. Histamine was purchased from Sigma.

### DNA constructs for expression in HeLa cells

Nir1-AA/RR-mCh was generated from Nir1-mCherry ^13^ by site-directed mutagenesis to replace Ala217 and Ala218 with Arginine residues. Nir2-AA/RR-mCh was generated from Nir2-mCherry ^7^ by site-directed mutagenesis to replace Ala516 and Ala517 with Arginine residues. Nir1-FLAG, Nir2-FLAG, and MAPPER constructs were described previously ^7,13^. GFP-PLCδ-PH was obtained from Dr. Tobias Meyer (Weill Cornell Medicine, New York, NY). The H1R plasmid was obtained from Dr. Elliott Ross (UT Southwestern Medical Center, Dallas, TX). The H1R-FLAG-IRES-GFP-PLCδ-PH construct was generated by assembling the H1R-FLAG fragment, and an internal ribosome entry site 2 (IRES2) sequence, and GFP-PLCδ-PH using In-Fusion^®^ multiple-insert cloning. H1R-FLAG and GFP-PLCδ-PH are separated by the IRES2 sequence obtained from Addgene plasmid #214425. All constructs were verified by either Oxford Nanopore sequencing (Plasmidsaurus Inc.) or Sanger sequencing (McDermott Center Sanger Sequencing Core, UT Southwestern Medical Center, Dallas, TX).

### Co-immunoprecipitation

HeLa cells were seeded in 6-well plates (CellStar, Cat. No. 657160) at a density of 1.0 x 10^6^ cells per well 24 h before transfection. Cells were co-transfected with FLAG- and mCherry-tagged constructs. After 16-20 h, cells were washed twice with ice-cold phosphate-buffered saline and lysed in Pierce IP Lysis Buffer (ThermoFisher; #87787) supplemented with 1X protease inhibitor cocktail (Sigma-Aldrich; P8340). Lysates were clarified by centrifugation at 12,000 × g for 30 min at 4°C. Supernatants were divided into input (40%) and immunoprecipitation (60%) fractions. The immunoprecipitation fraction was incubated overnight at 4°C with RFP-Trap magnetic affinity beads (Chromotek; rtma-20) in dilution buffer (10 mM Tris-HCl, pH 7.5, 150 mM NaCl, 0.5 mM EDTA). Beads were then separated using a magnet and washed three times with wash buffer (10 mM Tris-HCl pH 7.5, 150 mM NaCl, 0.05% Nonidet P-40, 0.5 mM EDTA). Bound proteins were eluted by resuspending beads in Laemmli sample buffer containing 10% β-mercaptoethanol and boiling at 100°C for 10 min. Input samples were treated identically. Samples were resolved on 7.5% or 10% Mini-PROTEAN TGX stain-free gels (Bio-Rad; #4568026 and #4568036) and transferred to polyvinylidene difluoride membranes. Membranes were blocked in 3% milk in Tris-buffered saline with 0.1% Tween-20 (TBST) for 1 h at room temperature, followed by overnight incubation at 4°C with primary antibodies (1:2000 mouse anti-FLAG, Sigma-Aldrich F1804; or 1:2000 mouse anti-mCherry, Abcam ab125096) in 4% milk TBST. Membranes were washed three times with TBST and incubated for for 1 h at room temperature with HRP-conjugated secondary antibody (1:2000 horse anti-mouse, Cell Signaling Technology #7076) in 4% milk TBST. After washing, signals were detected using Clarity Western ECL substrate (Bio-Rad; #170-5060) and visualized on film.

### Confocal and TIRF microscopy

HeLa cells were seeded on eight-well Nunc™ Lab-Tek™ chambered coverglass (Thermo Fisher Scientific) at a density of 1.5 x 10^4^ cells per well 24 h prior to transfection. After 16-20 h, cells were washed twice with extracellular buffer (ECB; 125 mM NaCl, 5 mM KCl, 1.5 mM MgCl_2_, 20 mM HEPES, 10 mM glucose, and 1.5 mM CaCl_2_, pH 7.4) and imaged in ECB. Confocal and total internal reflection fluorescence (TIRF) imaging were performed at room temperature using a custom spinning-disk confocal/TIRF imaging system built around a Nikon Eclipse Ti microscope. Images were acquired with either a 100× (NA 1.49) or 60× (NA 1.49) CFI Apochromat TIRF objective and captured with dual ORCA-Fusion BT sCMOS cameras. Illumination was provided by 405, 488, 561, and 640 nm solid-state lasers. Image acquisition and instrument control were carried out using Nikon NIS-Elements software.

### Nir2 translocation analysis using confocal microscopy

HeLa cells co-expressing Nir1-YFP and either Nir2-mCh or Nir2-AA/RR-mCh were stimulated with 100 µM histamine for 2 min. To quantify dynamic changes in Nir2 localization following stimulation, Nir1-YFP puncta were identified and defined as region of interest (ROIs) corresponding to ER-PM junctions. Fluorescence intensities of Nir2-mCh or Nir2-AA/RR-mCh within these ROIs were measured before and after stimulation, background-subtracted, and normalized to pre-stimulation levels.

### Nir2 translocation analysis using TIRF microscopy

HeLa cells co-expressing H1R, Nir1-YFP, and either Nir2-mCh or Nir2-AA/RR-mCh were stimulated with histamine at concentrations ranging from 100 pM to 10 µM for 1 min. Cells were imaged by TIRF microscopy to monitor changes in near-PM Nir2 fluorescence intensities. Images were acquired before and after stimulation. To quantify Nir2 recruitment, Nir1-YFP puncta were used to define region of interest (ROIs) corresponding to ER-PM junctions. For each cell, 19 puncta were randomly selected. A background ROI outside the cell was also measured. Fluorescence intensities of Nir2 within the ROIs were measured before and after stimulation, background-subtracted, and normalized to pre-stimulation levels.

### PIP_2_ replenishment assay

This assay was performed as previously described ^23^, with the modification that an H1R-FLAG-IRES-GFP-PLCδ-PH construct was used to ensure expression of H1R and GFP-PLCδ-PH at a fixed ratio in transfected cells. Relative fluorescence intensity of GFP-PLCδ-PH was monitored to quantify dynamic changes in PM PIP_2_ levels following receptor stimulation in HeLa cells transfected with H1R-FLAG-IRES-GFP-PLCδ-PH and the indicated Nir1 or Nir2 constructs. Single cells were imaged using TIRF microscopy. To minimize photobleaching, an ND8 neutral density filter was introduced into the light path during image acquisition. GFP-PLCδ-PH fluorescence intensity was measured over time, background-subtracted, and normalized to the initial time point.

## ACKNOWLEDGEMENTS

We thank the Liou and Airola Laboratory members for valuable discussions.

