## Supplemental Information for "Nir1-Nir2 Heterodimerization Expands the Sensitivity and Dynamic Range of the Phosphatidylinositol Cycle"

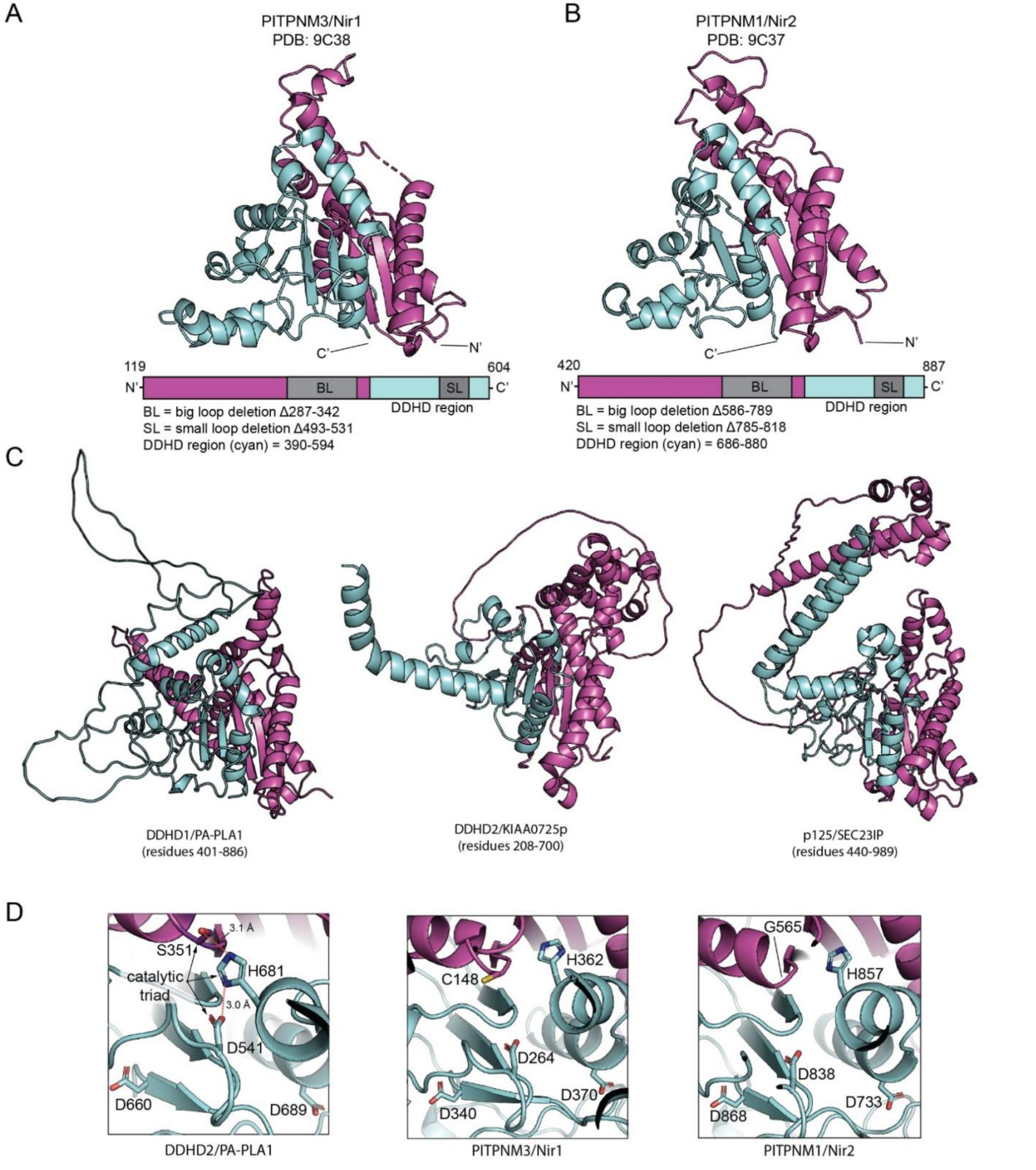


**Supplementary Fig. 1. DDHD region-containing proteins display the same domain topology as the Nir1 and Nir2 NirD experimental structures. (A, B)** The Nir1 (A) and Nir2 (B) NirD experimental structures highlighting the DDHD region (cyan) forms one half of the complete domain, with the other half formed by the preceding ~200 amino acids (purple). **(C)** AlphaFold2 predicted structures of the three PA-iPLA proteins present in humans (DDHD1, DDHD2, and p125) shown with their DDHD regions (cyan) combining with preceding amino acids (purple) to form a complete domain. **(D)** The triglyceride lipase, DDHD2, incorporates two DDHD-motif residues in its catalytic triad and AlphaFold places these residues in close proximity (left panel). Nir1 and Nir2 NirD domains have similarly placed DDHD-motif residues (middle and right panel) however they have an incomplete lipase motif that lacks an essential serine residue to complete the catalytic triad (GXSXG; S351 in DDHD2), and the Asp and His residues are located too far away to participate in hydrogen bonding.

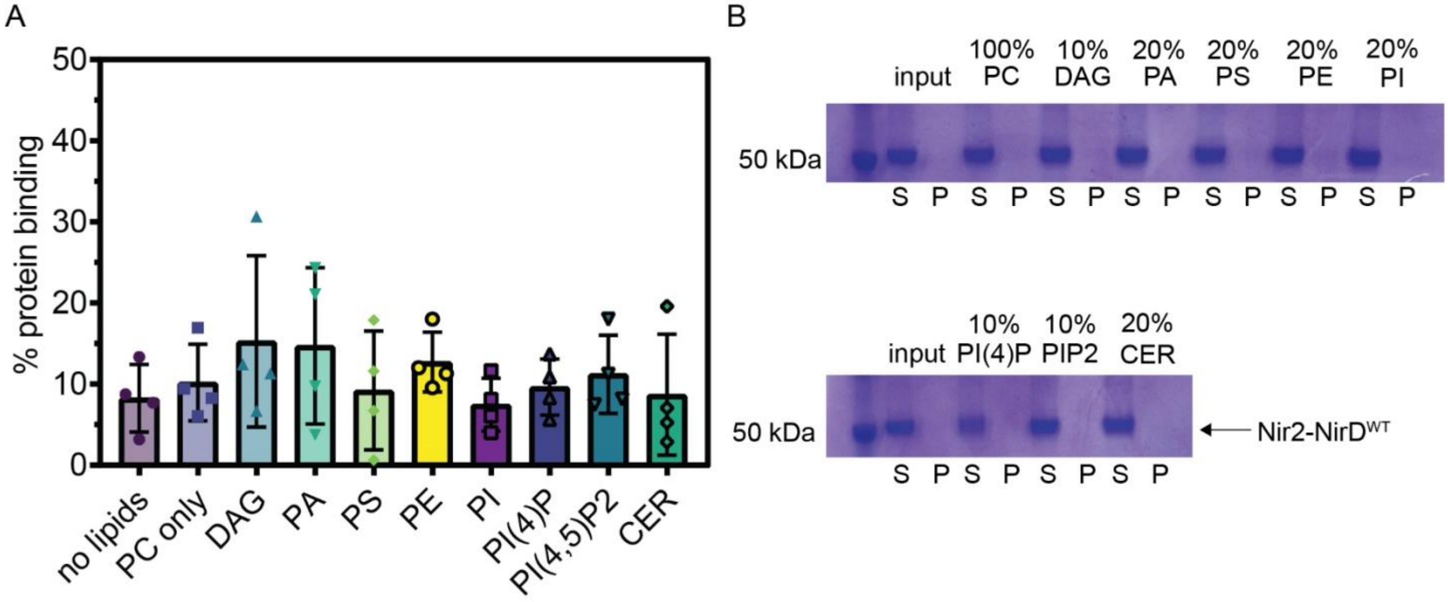


**Supplementary Fig. 2. The Nir2-NirD$^{WT}$ protein does not bind membranes. (A)** % protein binding to liposomes using a liposome co-sedimentation assay with purified Nir2-NirD$^{WT}$. The Nir2-NirD$^{WT}$ protein did not bind to liposomes under any lipid composition tested. The molar percentage of additional lipids were either 10 mol% (DAG, PI(4)P, PI(4,5)P2 or 20 mol% (PA, PS, PE, PI, Cer) in a PC liposome background. PC, phosphatidylcholine, DAG, diacylglycerol; PA, phosphatidic acid; PS, phosphatidylserine; PE, phosphatidylethanolamine; PI, phosphatidylinositol; PI(4)P, phosphatidylinositol-4-phosphate; PI(4,5)P2, phosphatidylinositol-4,5-bis-phosphate; CER, ceramide. **(B)** Representative SDS-PAGE gels of supernatant and pellet fractions for each liposome condition.

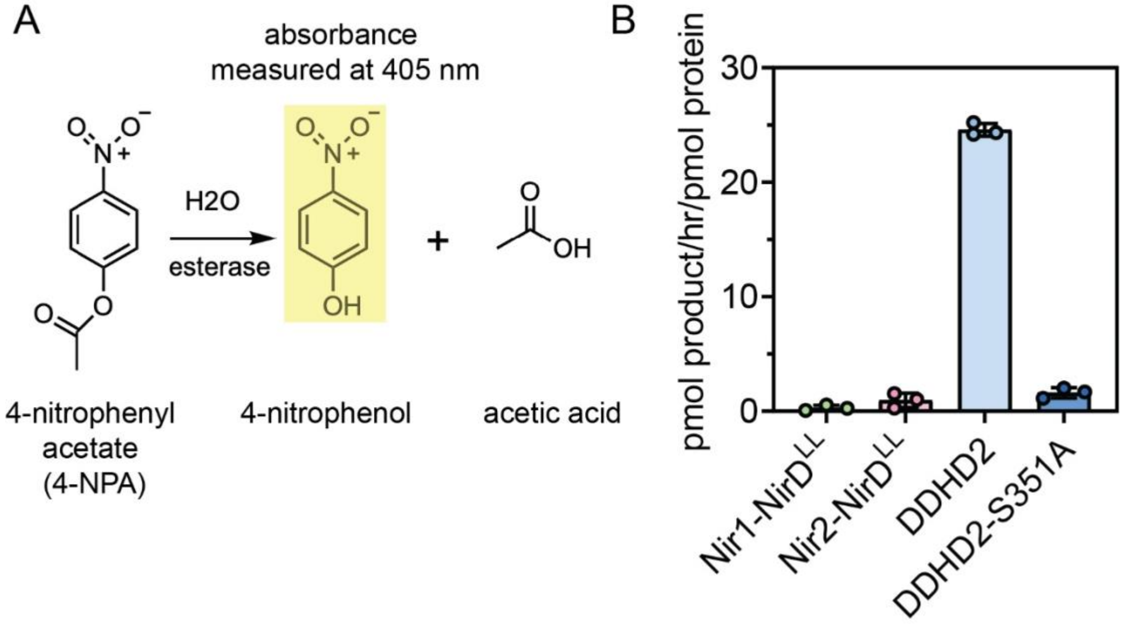


**Supplementary Fig. 3. The Nir1 and Nir2 NirD$^{LL}$ proteins do not exhibit detectable esterase activity against the generic esterase substrate 4-NPA. (A)** Schematic of the activity assay used to analyze the esterase activity of the Nir1 and Nir2 NirD$^{LL}$ domains, in comparison to a known esterase (DDHD2) and its inactive mutant (DDHD2-S351A). 4-NPA hydrolysis via esterase activity produces 4-nitrophenol (highlighted yellow) and its absorbance can be measured at 405 nm. The amount of product produced in the reaction can be equated to esterase activity. **(B)** Quantification of the amount of 4-nitrophenol produced in reactions of equal protein concentration in matching reaction buffer.

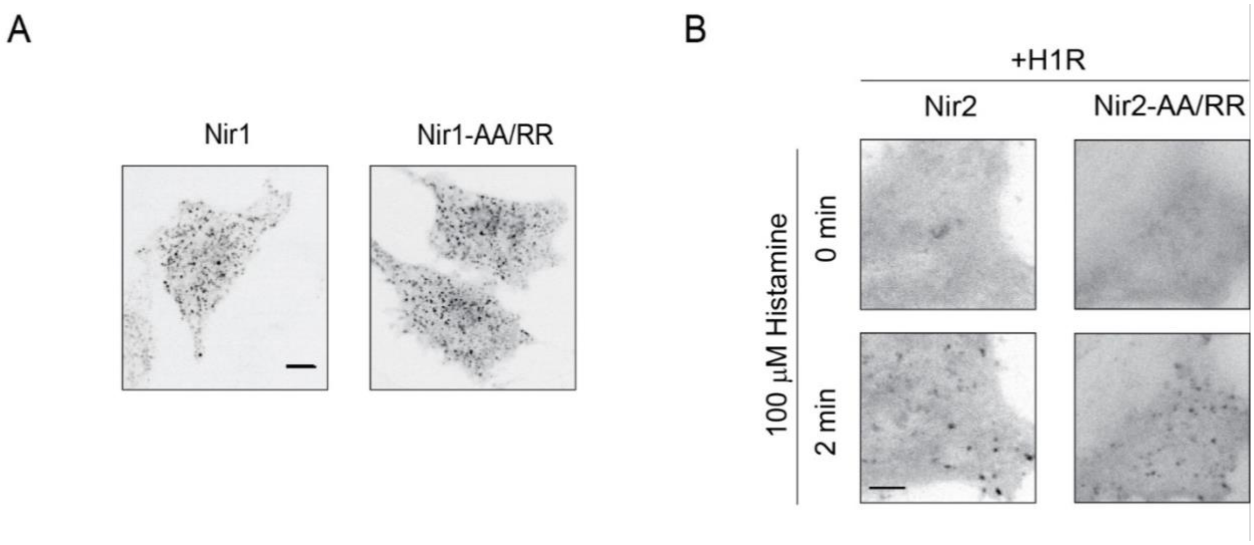


**Supplementary Fig. 4. AA/RR mutations do not affect the intrinsic abilities of Nir1 pre-localization or Nir2 translocation to ER-PM junctions. (A)** Confocal images of HeLa cells expressing Nir1-mCh or Nir1-AA/RR-mCh, showing comparable pre-localization to ER-PM junctions. Scale bar, 10 μm. **(B)** Confocal images of HeLa cells co-expressing the histamine H1 receptor (H1R) with either Nir2-mCh or Nir2-AA/RR-mCh following stimulation with 100 μM histamine for the indicated times. Scale bar, 10 μm.
